# Gamma oscillations during episodic memory processing reveal reversal of information flow between the hippocampus and prefrontal cortex

**DOI:** 10.1101/728469

**Authors:** Sarah Seger, Michael D. Rugg, Bradley C. Lega

**Affiliations:** Department of Neurological Surgery, University of Texas-Southwestern Medical Center, Dallas, Texas 75390; Center for Vital Longevity, University of Texas at Dallas, Dallas, Texas 75235; School of Behavioral and Brain Sciences, University of Texas at Dallas, Dallas, Texas 75080; Department of Psychiatry, University of Texas Southwestern Medical Center, Dallas, Texas 75390

## Abstract

A critical and emerging question in human episodic memory is how the hippocampus interacts with the prefrontal cortex during the encoding and retrieval of items and their contexts. In the present study, participants performed an episodic memory task (free recall) while intracranial electrodes were simultaneously inserted into the hippocampus and multiple prefrontal locations, allowing the quantification of relative onset times of gamma band activity in the cortex and the hippocampus in the same individual. We observed that in left anterior ventrolateral prefrontal cortex (aVLPFC) gamma band activity onset was significantly later than in the hippocampus during memory encoding, whereas its activity significantly preceded that in the hippocampus during memory retrieval. These findings provide direct evidence to support models of prefrontal-hippocampal interactions derived from studies of rodents, but suggest that in humans, it is the aVLPFC rather than medial prefrontal cortex that demonstrate these reciprocal interactions.

## Introduction

Prefrontal monitoring and control during episodic memory processing is thought to be critical for contextually mediated memory retrieval (***Miller, 2013***; ***Preston and Eichenbaum, 2013***). An influential model characterizing one of the mnemonic roles of the prefrontal cortex (PFC) – termed here the *reciprocal flow hypothesis* – posits that during memory encoding, contextual information flows from the hippocampus to the PFC while during retrieval, the PFC uses this stored information to guide selection of a contextually appropriate hippocampal memory representation (***Desimone and Duncan, 1995***; ***Desimone, 1998***; ***Miller and Cohen, 2001***; ***Preston and Eichenbaum, 2013***). Stated another way, the model posits that information flow between the hippocampus and PFC reverses direction between encoding and retrieval. Evidence supporting this model has come from rodent investigations employing lagged correlation between the hippocampus and PFC in theta band oscillatory power (e.g. ***Place et al., 2016***). In humans, noninvasive data have stimulated the hypothesis that the VLPFC is necessary for generating retrieval cues during episodic memory search (***Kim, 2019***), consistent with rodent findings, and lesion studies suggest that patients with frontal lobe dysfunction have difficulty recalling items when the context is altered between encoding and subsequent retrieval (***Chao, 1997***; ***Fletcher, 2001***). However, to date there is no direct human electrophysiological evidence of reversed lags in the timing of hippocampal and PFC activation that would be indicative of differential information flow during encoding and retrieval. fMRI studies lack sufficient temporal resolution to identify such an effect, precise source localization of MEG signals to different mesial temporal structures is problematic, and the absence of direct homology between rodent and human prefrontal cortex means that human intracranial EEG studies are necessary to establish whether this phenomenon is characteristic of human episodic memory and to determine in which brain regions it may occur.

A complicating factor when testing the reciprocal flow hypothesis in humans is that there appear to be multiple oscillations within traditional theta frequency bands, and the dominant theta frequency in the hippocampus may differ that in the neocortex (***Lega et al., 2012***; ***Miller, 2013***; ***Watrous and Ekstrom, 2014***). Furthermore, unlike rodents, human hippocampal recordings do not universally exhibit theta modulation as a function of memory processing (although this might be more prevalent in posterior hippocampal locations) (***Lin et al., 2017***; ***Watrous and Ekstrom, 2014***). By contrast, gamma oscillations exhibit widespread and reproducible power increases in multiple neural regions during episodic memory encoding and retrieval, including in the PFC and hippocampus (***Burke et al., 2014***; ***Sederberg et al., 2007***).

Here we sought evidence of reversal of information flow between the hippocampus and pre-frontal cortex during the encoding versus the retrieval of episodic memories. We did this by taking advantage of a unique dataset obtained from 77 human patients implanted with stereo EEG electrodes for seizure mapping purposes who performed a verbal free recall paradigm. During the study and recall phases of the task, we identified activation peaks in gamma oscillations from 40 to 120 Hz, using the onset of gamma activation as an estimate of the initial timing of activity in a given brain region. As our data set included subjects with electrodes implanted in both the hippocampus and PFC (in addition to other cortical locations), we were able to directly compare the timing of memory-related gamma activation in the PFC and hippocampus within-subjects.

## Results

### Behavioral Performance

Across participants, the average probability of recall for all words was 24.4%. The average percentage of list intrusions (recall errors) per subject was 12.8%. We derived an estimate of temporal clustering (the tendency for items adjacent to each other in the study list to be recalled sequentially) to determine if temporal contextual factors were operating at retrieval (***Watrous and Ekstrom, 2014***). The mean clustering factor across all participants was 0.642, robustly higher than the chance value of 0.500 (*t*(36) = 8.294, *p*< 0.001), indicating that participants incorporated temporal contextual information into encoded representations of the study words (***Sederberg et al., 2010***).

### sEEG Data

For our principal analysis, we identified the lag in onset of activation (Δ*t*_*γ*_) for five prefrontal locations relative to the hippocampus (positive Δ*t*_*γ*_ indicating activation following the hippocampus, negative Δ*t*_*γ*_ indicating activation preceding the hippocampus). We divided the PFC into five distinct bilateral regions: dorsal PFC (dorsal to the inferior frontal sulcus, anterior to pre-motor cortex, ventral to the superior frontal gyrus), the posterior VLPFC (posterior to the anterior ascending ramus of the sylvian fissure, anterior to motor cortex), the anterior VLPFC (anterior to that ascending ramus), the medial orbitofrontal cortex and the anterior cingulate cortex. These regions were selected based upon targeting strategies employed for seizure mapping, providing sufficient numbers of electrodes for analysis. Exact timing of activation onset (*t*_*γ*_) was estimated on a trial by trial basis for recording sites by calculating a gamma power threshold in the 40-120 Hz range to determine the timing of onset relative to hippocampal contacts *in the same individual* following established methods (Figure 1)

**Figure 1.**
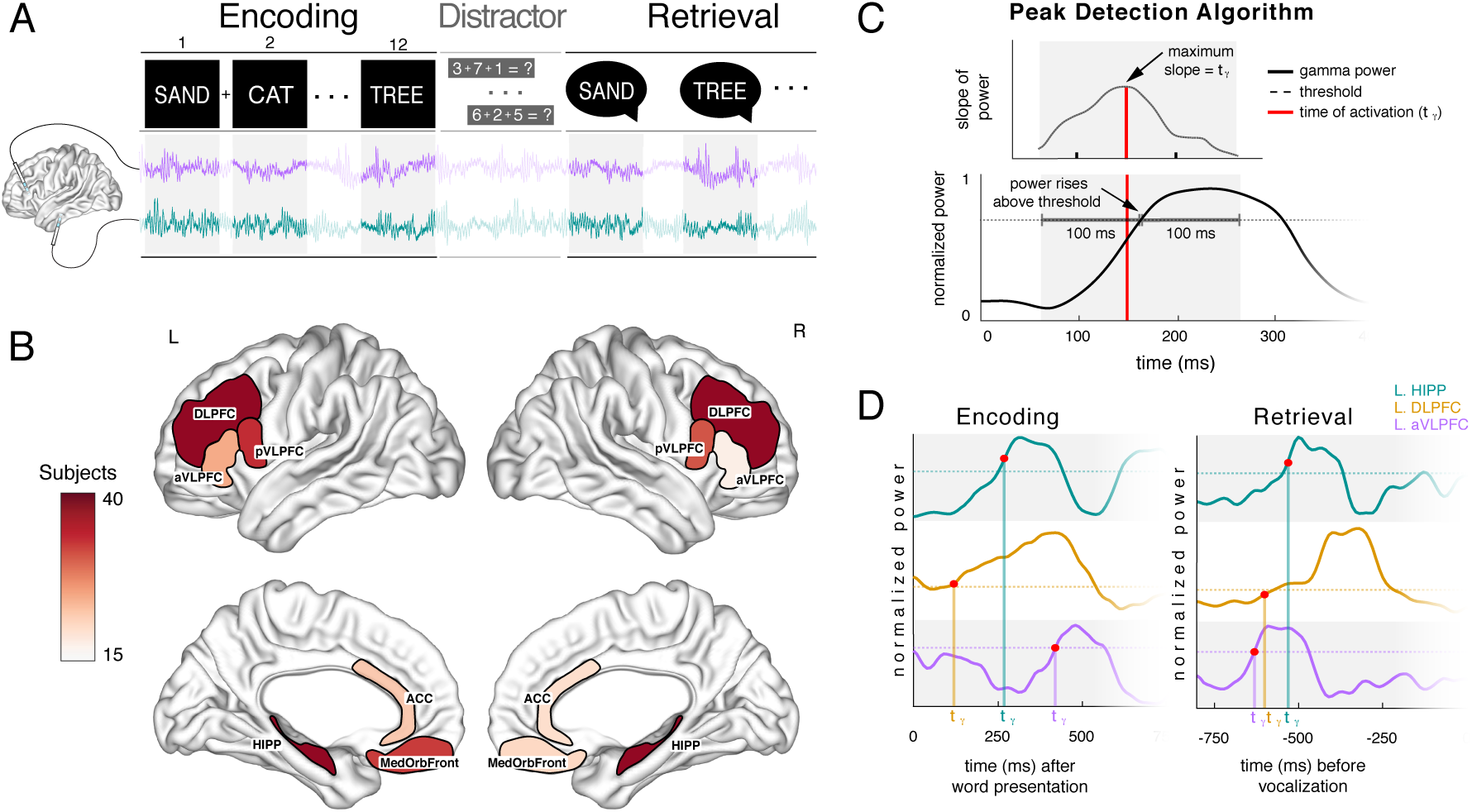
(A) Schematic of the experimental paradigm used in this study. Black boxes indicate the encoding and retrieval epochs. (B) Number of subjects included in each prefrontal and hippocampal region. The colorbar indicates number of subjects, a minimum of 15 subjects and a maximum of 40 subjects contributed to any given unilateral region. (C) Example trace from a single electrode contact depicts the activation onset detection algorithm. The time point of activation (*t*_*γ*_) is marked as the time point where the slope is maximized in the 200 msec window centered at the time point where power passes threshold. The *t*_*γ*_ demonstrates that onset of activation does not necessarily coincide with the time that power passes the threshold. (D) Example encoding and retrieval trial show *t*_*γ*_ for three electrodes. **Figure 1–Figure supplement 1**. **Figure 1–Figure supplement 2.**

**Figure 2.**
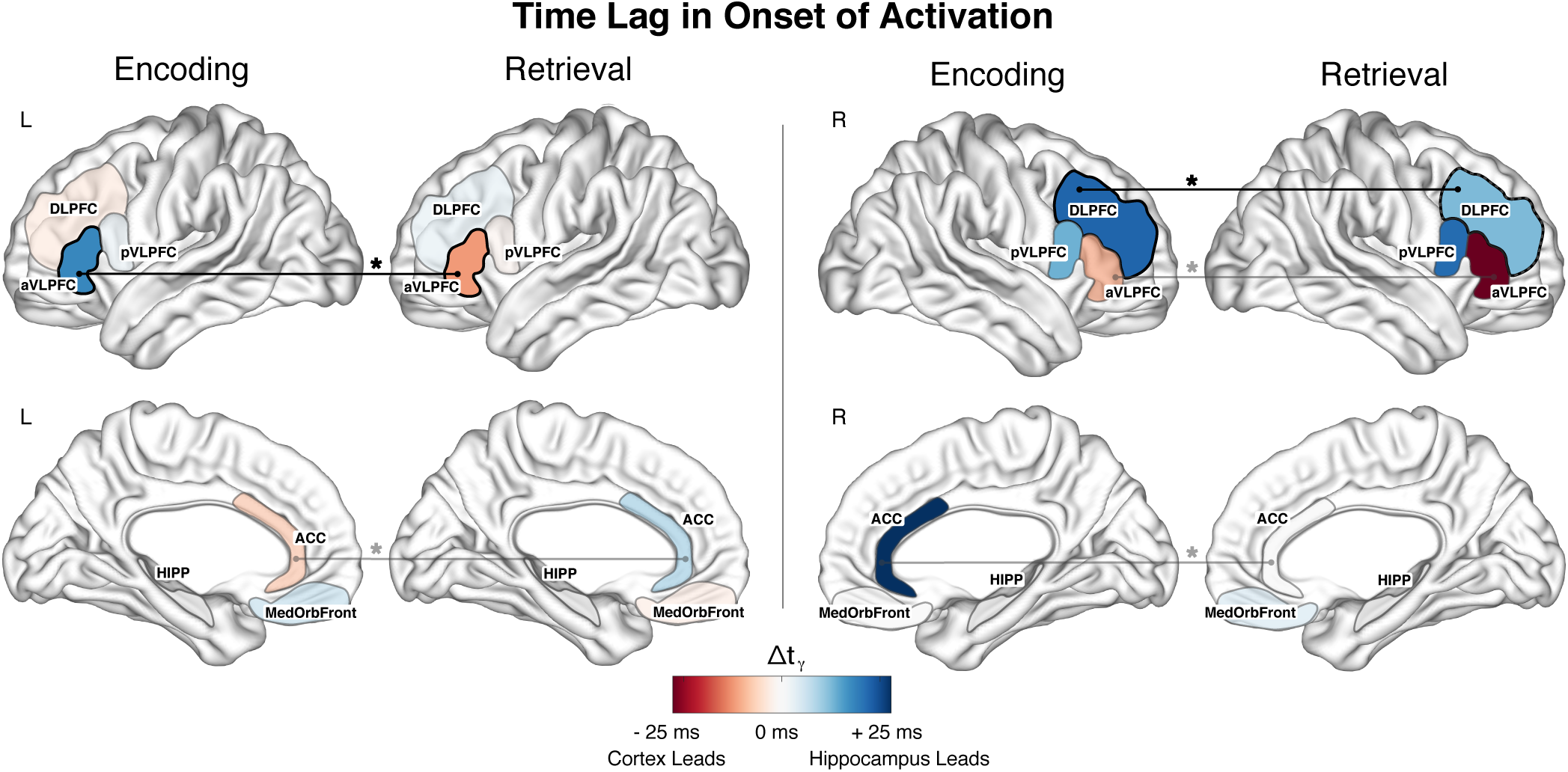
Mean Δ*t*_*γ*_ across electrodes for all prefrontal cortex regions for the encoding (recalled words only) and retrieval condition. Red colors indicate that activation in the hippocampus precedes activation in the cortex and blue indicates that the cortical activation precedes hippocampal. For each memory condition, a black region border indicates that Δ*t*_*γ*_ across all electrode pairs is significantly different than zero (t-test, FDR corrected *p*< 0.011 for encoding and *p*< 0.007 for retrieval). An asterisk (*) between the encoding and retrieval conditions indicates that the encoding and retrieval Δ*t*_*γ*_ are significantly different when compared with a paired t-test (FDR corrected *p*< 0.019). The left aVLPFC exhibited a mean activation lag relative to the hippocampus during successful encoding of +13.4 msec (FDR corrected *p*< 0.001, t-test of activation times compared to hippocampus), and a reversal of this effect during retrieval, such that the region led the hippocampus by −10.4 msec (FDR corrected *p* = 0.0116). Moreover, the Δ*t*_*γ*_ distributions for encoding and retrieval were significantly different (FDR corrected *p*< 0.001) across electrodes when compared with a paired t-test. Of the remaining PFC regions, only one other region, the right DLPFC, exhibited a Δ*t*_*γ*_ that was significantly greater than zero during encoding (FDR corrected *p* = 0.0037); however this region did not exhibit a reversal in Δ*t*_*γ*_ values during retrieval (with the hippocampus leading during both encoding and retrieval). The left ACC exhibited a significant difference in the distribution of Δ*t*_*γ*_ during encoding vs. retrieval (−6.86 msec during encoding, 6.46 msec during retrieval; FDR corrected *p* = 0.0173); but the Δ*t*_*γ*_ was not significantly different than zero during neither encoding (FDR corrected *p* = 0.0700) or retrieval (FDR corrected *p* = 0.0700) indicating onset nearly commensurate with that of the hippocampus. The pattern of Δ*t*_*γ*_ reversal was evident for the left but not the right aVLPFC. In the latter region, while there was a significant difference between encoding and retrieval (FDR corrected *p* = 0.0080), the values indicated that the cortex led the hippocampus during both phases of the free recall task (lag = −23.5 msec during retrieval, −7.44 msec during encoding). **Figure 2–Figure supplement 1.**

Convincing evidence of a reversal in the flow of information consistent with the reciprocal flow model would require a PFC region to exhibit 1) a lag in activation onset relative to the hippocampus that is significantly greater than zero during successful item encoding (significant positive Δ*t*_*γ*_, hippocampus leading), 2) a lag in activation that is significantly less than zero during retrieval (negative Δ*t*_*γ*_, hippocampus trailing), and finally 3) a significant difference in these Δ*t*_*γ*_ values when directly compared in a paired test (reversed Δ*t*_*γ*_). Across all electrodes in our dataset (without any filtering of electrodes based upon their functional properties), we observed that the left aVLPFC exhibited the following set of properties: a mean activation lag relative to the hippocampus during successful encoding of 13.4 msec (FDR corrected *p* < 0.001, t-test of activation times compared to hippocampus), and a reversal of this effect during retrieval, such that the region led the hippocampus by −10.4 msec (FDR corrected *p* = 0.0116). Moreover, the Δ*t*_*γ*_ distributions for encoding and retrieval were significantly different (FDR corrected *p* < 0.001) (Figure 1).

Next, we compared the Δ*t*_*γ*_ distributions during successful versus unsuccessful encoding, looking for evidence of a subsequent memory effect in this measurement. For the left aVLPFC, this contrast was significant (FDR corrected *p* = 0.0216), suggesting that Δ*t*_*γ*_ measurements are sensitive to encoding success (Figure 3). Taken together, these findings indicate that the timing of gamma activation onset in the left aVLPFC constitute a signal that is sensitive to memory encoding success (exhibiting an SME), with a pattern that fits a putative model of the transfer of contextual information to the frontal cortex during successful encoding (significantly positive relative to hippocampal activation) with evidence of a reversal during retrieval (significantly negative relative to the hippocampus). Across all electrode pairs, 38% of aVLPFC electrodes exhibited this pattern of Δ*t*_*γ*_ reversal, which was significantly greater than the fraction exhibiting this effect in the DLPFC (*χ*^2^(1, N=630) = 17.160, *p* < 0.001) (Figure 4). Across the subjects who contributed an electrode pair to the left aVLPFC, 59% showed a pattern of Δ*t*_*γ*_ reversal in at least one electrode pair.

**Figure 3.**
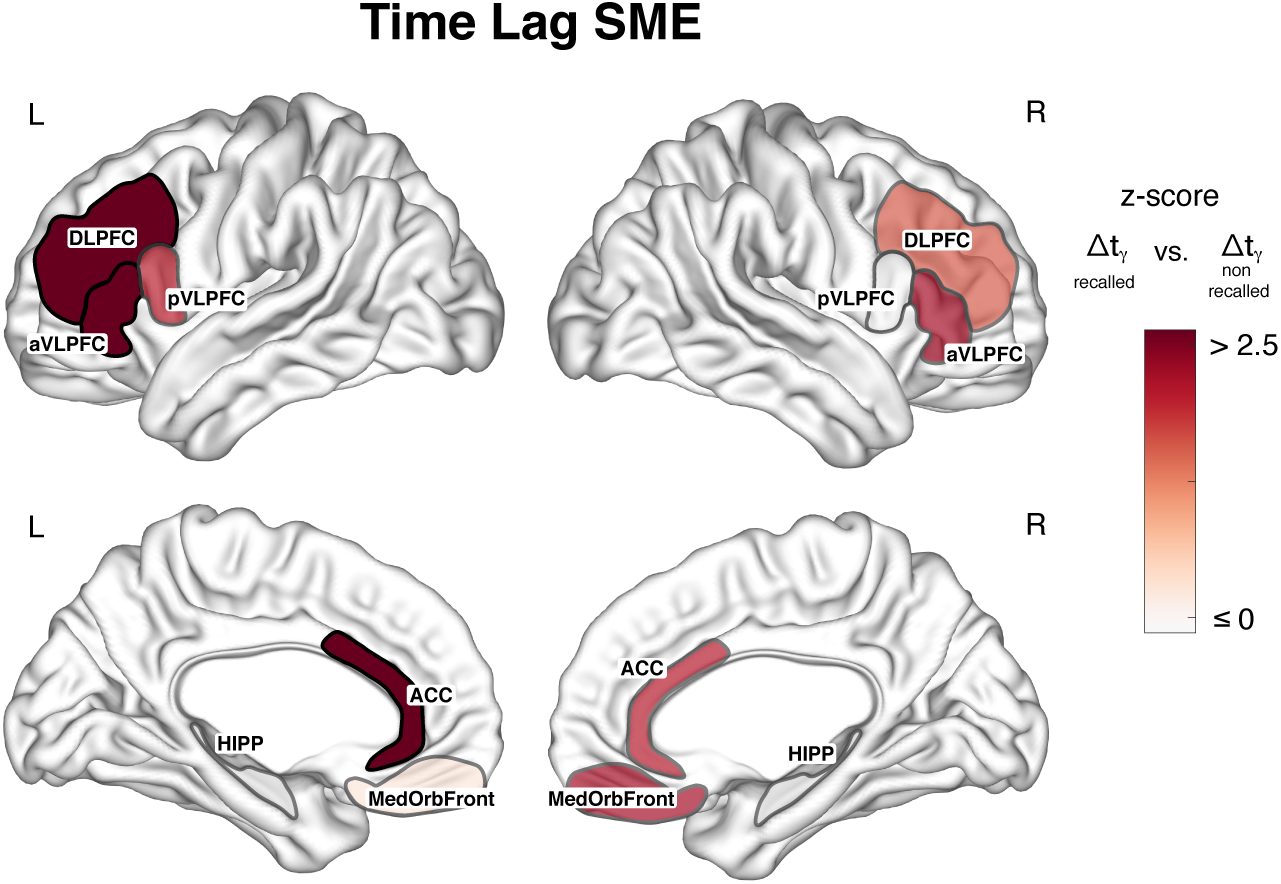
Subsequent memory effect in Δ*t*_*γ*_ for all regions. Z-scores were calculated for each region using a paired t-test between Δ*t*_*γ*_ for recalled words and Δ*t*_*γ*_ for non-recalled words. The subsequent memory effect for left aVLPFC, left DLPFC, and left ACC was significant (FDR correctedp< 0.007), which is indicated by black borders for those regions. No regions in the right hemisphere show a significant subsequent memory effect. **Figure 3–Figure supplement 1.**

**Figure 4.**
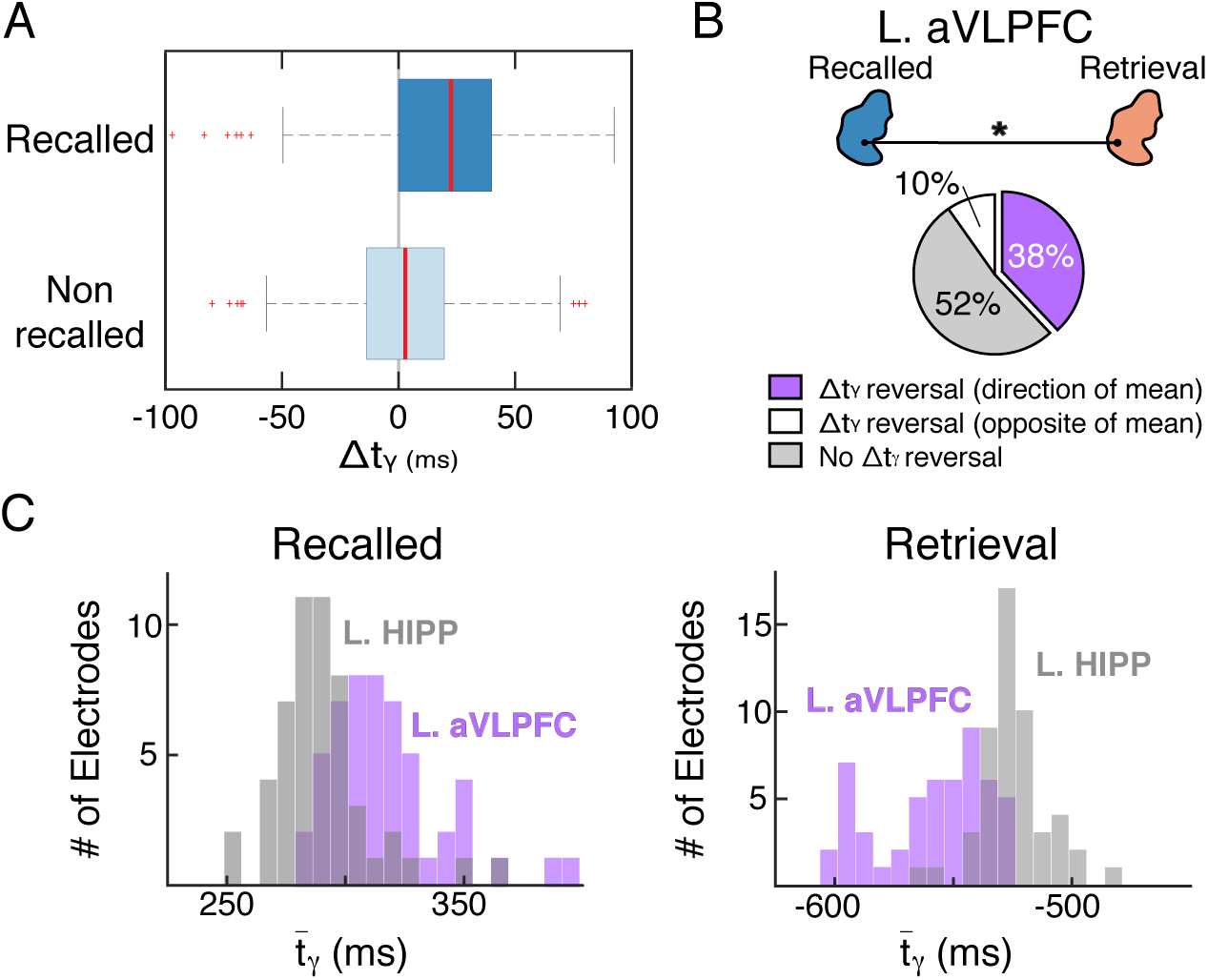
The left aVLPFC shows a reversal in the Δ*t*_*γ*_ between encoding and retrieval consistent with the reciprocal flow information. (A) Distribution of Δ*t*_*γ*_ for all aVLPFC electrodes during the recalled and non-recalled conditions. The Δ*t*_*γ*_ for non-recalled words is not significantly different than zero (mean non-recalled lag is +2.91 msec). (B) 38% of aVLPFC electrodes have Δ*t*_*γ*_ reversal between conditions, with the hippocampus leading in activation during encoding and the cortex leading during retrieval, and 10% show the opposite pattern of activation, with the cortex leading in activation during encoding and the hippocampus leading during retrieval. 52% of electrodes show no reversal in lag between conditions. (C) Histograms of the mean 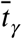 during encoding (subsequently recalled only) and retrieval for the 38% of aVLPFC – HIPP electrode pairs exhibiting the effect depicts the differences in timing of activation onset for hippocampal and aVLPFC electrodes between memory conditions.

Further, we analyzed prior list intrusions (PLI) to test more directly whether Δ*t*_*γ*_ reversal is associated with the transmission of contextual information, as hypothesized by the reciprocal flow model. List intrusions represent errors of item–context association (the wrong item for a given context, i.e. the list on which the item was presented, although we discuss caveats to the interpretation of PLI data in the Discussion below). For this analysis, oscillatory activity can be analyzed only during item retrieval. We observed no evidence of information reversal for PLI events, with the onset of left aVLPFC activation not significantly different than for the hippocampus (mean Δ*t*_*γ*_ = −3.5 msec, uncorrected *p* = 0.3861). In addition, the Δ*t*_*γ*_ during correct retrieval events was significantly less than that of PLI events (uncorrected *p* = 0.0436).

In a convergent approach, we looked for evidence of lagged activation using a different method, this time employing the lagged correlation of the gamma power envelope (rather than activation onset), following established methods (***Ossandon et al., 2011***). With this approach, which has a very different rationale and underlying set of assumptions than the lagged activation analysis described above, the results for the left aVLPFC were highly consistent with those observed previously, with a significant positive lag of 14 msec during encoding (uncorrected *p* = 0.0442) (hippocampus leading) and negative 8 msec lag during retrieval (in the same direction as our initial analysis with the PFC leading, though it did not reach significance – uncorrected *p* = 0.1426). As with the previous method, the encoding versus retrieval lag distributions were significantly different (FDR corrected *p* = 0.0276).

## Discussion

Our data reveal direct human electrophysiological evidence of the reversal of information flow between the hippocampus and prefrontal cortex (left aVLPFC) during episodic memory encoding versus retrieval. These results are analogous to observations in rodents held to be consistent with the reciprocal flow model, by which contextual information is transmitted to the prefrontal cortex during item encoding and then utilized to guide selection of an appropriate memory representation during retrieval (see ***Preston and Eichenbaum, 2013***). Our findings indicate that the left aVLPFC (but not pVLPFC, DLPFC, OFC, or anterior cingulate) exhibit the specific combination of characteristics (positive lag in activation onset during encoding, negative lag during retrieval) that one would predict based upon the reciprocal flow model. The specificity of the effect suggests this is a focal phenomenon in the prefrontal cortex in humans. Of interest, fMRI, BOLD SMEs are consistently reported for the aVLPFC and surrounding regions in study tasks requiring elaborative encoding of verbal items (for a review, see ***Kim, 2019***).

Using data drawn from a series of verbal memory tasks, Badre et al. proposed that the aVLPFC supports cue specification (***Badre et al., 2005***), and Simons and Spiers integrated similar previous findings into a model that distinguished between ventral and dorsal PFC contributions to verbal memory encoding and retrieval (***Simons and Spiers, 2003***), with ventral regions providing cue specification consistent with onset of activation preceding the hippocampus during retrieval, as we observed. In the setting of the free recall paradigm, cue specification presumably includes specification of temporal contextual information (***Sederberg et al., 2010***). We acknowledge however that our data by itself does not allow us to make strong claims regarding the content of information characterized by reciprocal flow.

Since we did not employ an experimental manipulation that allow a distinction to be made between verbal information and its contextual features, it is possible that our observations could be consistent with a contribution from the aVLPFC at encoding that is specifically related to the verbal features of the items such as cue processing (e.g. selecting a specific meaning for an item), at retrieval, the transmission of a non-specific ‘biasing’ signal that favors a hippocampal “retrieval mode” (e.g. ***Lepage et al., 2000***).

That being said, in the rodent investigations that motivated our analysis, similar observations to ours were interpreted as evidence of the transmission of contextual information, and the lack of a significant reversal in aVLPFC during retrieval of list intrusions is consistent with the proposal that lag reversal in this region is related to the transmission specifically of temporal contextual information. We acknowledge however that this inference is weakened by the concern that list intrusions for freely recalled items, may occur for a variety of reasons other than the failure to specify the correct list context, such as false binding of an item to the wrong temporal context, weak memory, and strong yet acontextual memories (strong familiarity) (***Ranganath, 2010***; ***Sederberg et al., 2008***; ***Yonelinas, 1999***). As it is not clear a priori what predictions the reciprocal flow model would make for each of these situations, definitively linking our observations to the transmission of context information may require an alternative memory paradigm, such as context switching, as has been implemented to demonstrate memory errors in patients with prefrontal lesions (***Badre and Wagner, 2007***; ***Rossi et al., 2009***). Nonetheless, previous rodent and human studies have provided evidence for the relevance of prefrontal regions in supporting context representations (***Desimone and Duncan, 1995***; ***Desimone, 1998***; ***Miller and Cohen, 2001***).

Rodent studies of prefrontal activation during contextually mediated mnemonic processing have mostly focused on the rodent medial PFC (mPFC) (***McKenzie et al., 2016***), whereas we observed evidence consistent with information reversal in the lateral cortex. However, it is an understatement to say that the homology between primate and rodent prefrontal regions is not fully understood and is likely limited (***Passingham et al., 2012***). Our dataset did not include strong representation of mPFC regions in terms of electrode coverage apart from the anterior cingulate. In this region, the electrode distribution may not have encompassed the cingulate regions observed to exhibit context-related BOLD activation in human noninvasive studies (***King et al., 2015***; ***Anderson et al., 2010***). This focus on the mPFC is motivated by direct anatomical connectivity between the anterior cingulate cortex and the hippocampus in humans (along with indirect connections between medial Brodmann area 10 and the hippocampus) (***Carmichael and Price, 1995***). These data have been interpreted as evidence that the mPFC supports the integration of specific items into contextual/semantic “schemas” (***Preston and Eichenbaum, 2013***; ***Schlichting and Preston, 2015***). It will prove insightful to directly compare PFC-hippocampal activation timing between aVLPFC and mPFC regions in an expanded dataset, looking for how this hypothesis may complement reversal of information.

Our principal results identifying Δ*t*_*γ*_ reversal (consistent with rodent findings) were obtained using an unbiased approach incorporating all electrodes in our dataset, but we acknowledge that the magnitude of the Δ*t*_*γ*_ offset depends upon whether one includes all electrodes in the calculation or only those that exhibit the reversal (a minority of electrodes in the aVLPFC exhibited the opposite pattern to the overall effect, and therefore, opposite to that predicted by the reciprocal information flow hypothesis). When estimated from the 38% of aVLPFC electrodes that exhibit encoding/retrieval reversal in Δ*t*_*γ*_, the mean encoding Δ*t*_*γ*_ is +29.3 msec and the mean retrieval Δ*t*_*γ*_ is −43.1 msec. Therefore, we do not make any claims about the relationship between Δ*t*_*γ*_ reversal and the precise offset of timing relative to gamma or theta cycles, unlike in rodent investigations (e.g. ***Place et al., 2016***). Taken together, our data support the relevance of the reciprocal flow hypothesis to human memory and establish lagged gamma activation as a method to identify functional interactions between memory-relevant regions in humans. The identification of electrode contacts in the aVLPFC that exhibit these functional properties may be a strategy for the identification of propitious targets of neuromodulation to treat memory disorders.

## Methods and Materials

### Participants

77 patients with medically intractable epilepsy who underwent stereoelectroencephalography surgery for clinical purposes were recruited to participate in this study. In total, there were 46 men and 31 women between 21-64 years of age. Data were collected from the University of Texas Southwestern Medical Center epilepsy program over a period of 3 years. Electrode placement was dictated solely by the clinical need for seizure localization. Each subject was implanted with up to 17 depth electrodes containing 8-16 cylindrical platinum–iridium recording contacts spaced 2-6-mm apart. Following implantation, electrode localization was achieved by co-registration of the post-operative computed tomography scans with the pre-operative magnetic resonance images using the FSL’s/*FLIRT* software (https://fsl.fmrib.ox.ac.uk/fsl/fslwiki/FLIRT). The co-registered images were evaluated by a member of the neuroradiology team to determine the final electrode locations. Each subject provided simultaneous recordings from the hippocampus and the frontal cortex. Frontal contacts were divided into 5 regions: the anterior ventro-lateral prefrontal cortex (aVLPFC) (principally BA45), the posterior ventro-lateral prefrontal cortex (pVLPFC) (BA8, BA44), the dorsolateral prefrontal cortex (DLPFC) (BA9, BA46), the medial orbitofrontal cortex (MedOrbFront) (BA10, BA11, BA12), and the anterior cingulate cortex (ACC) (BA24). The superior frontal sulcus was used to define the limits of the DLPFC dorsally, and the inferior frontal sulcus defined its ventral extent. The VLPFC was divided into anterior and posterior regions relative to the ascending ramus of the sylvian fissure. Orbitofrontal contacts were anterior to the anterior ramus and medial to the lateral orbital sulcus. Cingulate electrodes were inserted using the cingulate sulcus as a guide; all electrodes were ventral to the dorsal limit of the genu of the corpus callosum. Brodmann areas are provided as estimates of localization. The anatomical features described above were used in expert neuroradiology review to localize all electrodes and in situations of conflict between the reviewed anatomical location and Talairach-based assignment to Brodmann areas the former was used for localization. Electrodes corresponding to the site of ictal onset were excluded from analyses by direct expert neurology review (13 electrodes from 9 subjects).

### Experimental Paradigm

Participants preformed a free recall task consisting of multiple study/test cycles. During the study period, 12 words from a pre-selected pool of high-frequency, single-syllable, common nouns were visually presented, one at a time, on a computer screen for a duration of 1.6 s followed by a blank screen of 4 s with 100 msec of random jitter. Subjects were instructed to study each word as it appeared on the screen. The presentation of the last item in a list was followed by a 30 s period during which a math distractor task (A + B + C = ??) was performed to limit rehearsal. Participants were then instructed to verbally recall as many items as possible from the immediately prior list in no particular order. A full session consists of 12 full study/test cycles and 1 practice study/test cycle which was excluded from analysis. One complete session yielded electrophysiological recordings from 144 word encoding epochs (12 lists × 12 words) and a variable number of retrieval epochs. Participants performed between 1 and 9 sessions of the free-recall task over several days (median number 2).

We used the temporal clustering factor, which is a measure of temporal contiguity for each recall transition relative to all possible recall transitions at a given time, to determine if contextual factors were operating at retrieval (***Sederberg et al., 2008***). A temporal factor of 1 indicates a transition to the most temporally proximate item, whereas a temporal factor of 0 indicates a transition to the least temporally proximate item and 0.5 indicates a random transition. The temporal factor was averaged across all transitions to obtain a single estimate of temporal contiguity for each subject.

### Data Processing

Stereo-EEG data were recorded using a Nihon Kohden EEG-1200 clinical system. Signals were sampled at 1000 Hz and referenced to a common intracranial contact. Raw signals were subsequently re-referenced to a bipolar montage, with each contact referenced to the superficial adjacent contact. All analyses were conducted using MATLAB with both built-in and custom-made scripts. We employed an automated artifact rejection algorithm to exclude interictal activity and abnormal trials (kurtosis threshold greater than 4). The raw signals were filtered for noise on a session by session basis using the following steps: 1) the power spectral density was estimated across the entire session, 2) a 7th order polynomial was fit to the power spectral density estimate to obtain a trend line, 3) the trend line was subtracted from the power spectral density estimate to identify peaks in the periodogram, and 4) for each peak above 15 dB, the local minima surrounding the peak were used to define the cutoff frequencies for a first-order Butterworth notch filter. The notch filter identified for each peak in the periodogram was applied sequentially to the raw data. Retrieval trials were isolated such that each included trial was isolated from any other retrieval events by at least 1200 msec before the onset of vocalization and 200 msec after the onset of vocalization (this led to the exclusion of 3,258/12,791 [25.5% trials]). Identification of retrieval events followed previously published methods (***Burke et al., 2014***).

To assign bipolar electrode contacts to regions of interest, electrodes were defined as being in the hippocampus or one of the five PFC regions if at least 1 of the bipolar contacts was determined to lie within the structure. To compare activation onset times between electrodes in the hippocampus and the PFC for a given subject, each electrode contact within a given PFC region was paired with all hippocampal contacts for that subject (PFC-hippocampal electrode pairs). This was repeated for all regions.

### Activation Onset Detection (Calculation of *t*_*γ*_)

We compared the temporal patterns of high gamma band power changes in the hippocampus and frontal cortex in the 1000 msec immediately following study item presentation (encoding) and the 1000 msec immediately preceding word vocalization (retrieval). The bipolar sEEG from each encoding and retrieval epoch along with a 12 s flanking buffer was first bandpass filtered between 40 and 118 Hz (Barlett-Hanning, 1000th order) to reduce any possible influence of lower frequencies (***Kucewicz et al., 2019***) and then notch filtered from 59 to 61 Hz (Barlett-Hanning, 1000th order) to reduce possible line noise and then subjected to spectral decomposition into 10 msec time bins and 20 frequency bands from 40 to 120 Hz using a multi-taper fast Fourier transform (taper parameters: 3 tapers, time-bandwidth product of 2, 200 msec moving window, 10 msec step size) (Chronux toolbox, RRID:SCR_005547). The decomposed spectral power values within each frequency band were z-scored separately for all encoding and retrieval epochs in a given session by subtracting the mean and dividing by the standard deviation to provide a power spectrum that characterized oscillatory activity at each site separately for memory encoding and memory retrieval. Normalized power estimates were averaged into 4 gamma bands to obtain an overall estimate of high gamma activity (40-50 Hz, 50-60 Hz, 60-80 Hz, and 80-118 Hz) (Figure 1C,D).

To obtain an estimate of gamma power, a threshold was defined as the mean spectral power within each frequency band across all trials for a given session and for each condition. Prior to obtaining the threshold for encoding trials, the trials were further divided into subsequently recalled and non-recalled items to account for possible power differences due to memory success. For each encoding and retrieval trial, if the gamma power trace remained above a value of 1.5 times the calculated threshold for a minimum of 100 msec (approximately 3 cycles of the gamma band lower cutoff frequency) then it was included for analysis (***Dastjerdi et al., 2011***). On average, 98.1% of recalled, 98.3% of non-recalled, and 95.5% of retrieval trials were included for analysis. For each trial that passed the threshold criteria, a 200 msec window, centered at the time point where power first rose above threshold, was divided into 20 msec non-overlapping time bins, the slope was calculated as the difference in power between the two time points in the bin, and the start time of the bin with the largest slope was defined as the onset time (*t*_*γ*_) of high gamma activity for that trial (***Dastjerdi et al., 2011***). This was repeated for each frequency band and trial, and the *t*_*γ*_ was averaged across all bands and then across all trials for the recalled epochs, non-recalled epochs, and retrieval epochs to obtain a single time estimate of activation onset, 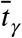, for each hippocampal and PFC electrode and each condition (see Figure 2). We repeated our analysis using a threshold of 2 times the calculated threshold, without a change in results for the aVLPFC (see Figure 2-Figure supplement 1).

To compare the onset time 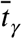 of gamma power activity (described above) of hippocampal electrodes to those in the PFC, a lag value, Δ*t*_*γ*_, of the gamma power activation onset was calculated for each PFC-hippocampal electrode pair and defined as

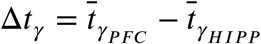

where 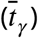 is the estimate of activation for the PFC and hippocampus, respectively. Given this, a positive Δ*t*_*γ*_ indicates that activation in the hippocampus is *leading* activation in the PFC, whereas negative Δ*t*_*γ*_ indicates that activation in the hippocampus is *lagging* activation in the PFC. For each hippocampal-PFC electrode pair, the Δ*t*_*γ*_ was calculated separately for the encoding (recalled and non-recalled separately) and retrieval trials to yield a single Δ*t*_*γ*_ per electrode pair for each condition.

### Cross-Correlation

In a convergent analysis, we used cross-correlation of the gamma band amplitude envelope to investigate the temporal dynamics between the hippocampus and the PFC. Trials were again partitioned into encoding (recalled and non-recalled) epochs, defined here as a 2600 msec window starting 500 msec prior to word presentation, and retrieval epochs, defined as a 2000 msec window starting 1500 msec prior to word vocalization. The bipolar sEEG for each epoch was bandpass filtered between 40 and 58 Hz (Barlett-Hanning, 1000th order), with the upper cutoff frequency selected to minimize contamination from line noise. The filtered signal was then Hilbert transformed and squared, from which the DC component (mean) across each epoch and frequency band was subtracted to obtain the mean-centered instantaneous amplitude envelope. For each hippocampal-PFC electrode pair, the normalized cross-correlation was calculated on the mean-centered amplitude envelope on a trial-by-trial basis using an 800 msec moving window with a 1 msec step size and a maximum lag of 150 msec. The initial 800 msec cross-correlation moving window for the encoding epochs was centered 100 msec prior to word presentation and stepped by 1 msec until 1700 msec after word presentation. The initial retrieval cross-correlation was centered 1100 msec prior to word vocalization and stepped by 1 msec until 100 msec after word vocalization. Thus the resulting matrices for the cross-correlations of a single trial were 301 by 1800 for encoding and a 301 by 1200 for a retrieval. For each electrode pair, the correlation coefficients were Fisher transformed across the recalled, non-recalled, and retrieval trials separately to allow comparison across trials (***Cohen et al., 2003***). Next for each hippocampal-PFC electrode pair and each condition, the Fischer transformed correlation coefficients for each correlation window and each correlation lag were tested against zero across trials using a one-sample t-test to generate a single matrix of p-values for each condition and each electrode pair. The resulting p-values were then normalized with an inverse transformation to allow for comparison across correlation lags and time points. Lastly, the correlation lag time with the highest z-score (i.e. the correlation lag where the correlation coefficient was maximally greater than zero across trials) was determined for each correlation moving window to produce a 1 by 1800 matrix for all recalled and non-recalled study trials and a 1 by 1200 matrix for all retrieval trials for each electrode pair.

### Statistical Procedure

To test for significant Δ*t*_*γ*_ across the electrodes within each hippocampal-PFC region pair during the encoding (subsequently recalled only) and retrieval conditions, we combined the Δ*t*_*γ*_ for all electrode pairs into a single matrix and used a t-test to compare the distribution of Δ*t*_*γ*_ against a null hypothesis of zero lag in onset activation (Δ*t*_*γ*_ = 0). In order to account for the type I error rate, p-values were false discovery rate (FDR) corrected. To test for differences in Δ*t*_*γ*_ between recalled/non-recalled and the recalled/retrieval conditions, we used a two-sample paired t-test with FDR correction.

For the cross-correlation analysis, we averaged the correlation lag at which the correlation coefficient was maximally greater than zero across all correlation windows for each electrode pair and each condition in order to obtain a single estimate of direction per electrode pair and condition. A one-sample t-test was used to compare the distribution of lag estimates across electrode pairs against a null hypothesis of zero lag (i.e. the correlation coefficient is maximized at zero lag) for each condition. A two-sample, paired t-test was used to compare the distribution of lag estimates for the encoding and retrieval conditions across electrodes against a null hypothesis of no difference in lag between conditions.

---

**Figure 1–Figure supplement 1.**
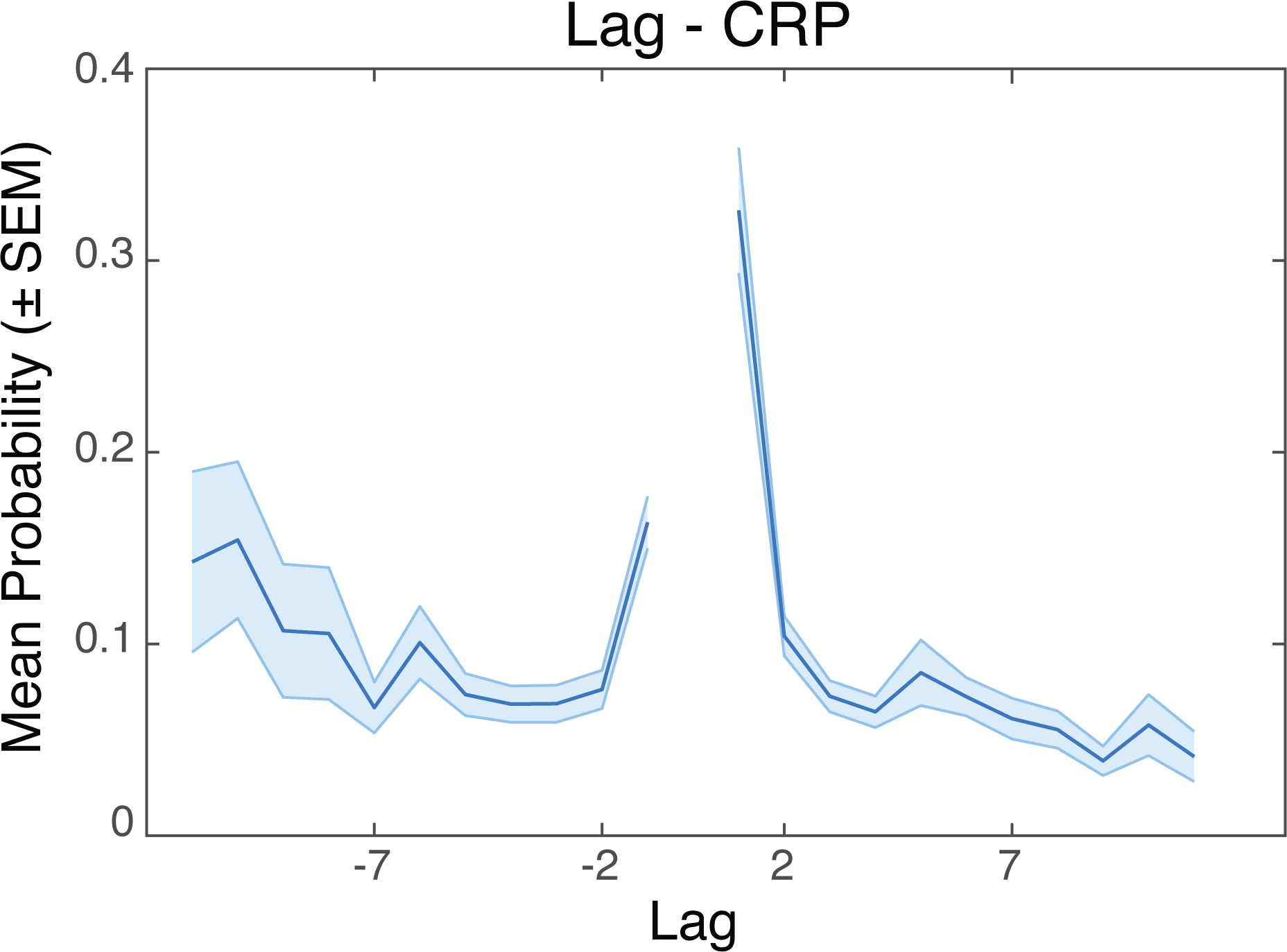
Average conditional response probability as a function of serial position lag. Error bars represent 95% confidence intervals across all subjects. Higher probabilities for the lags closer to zero demonstrate a tendency for items adjacent to each other in the study list to be recalled sequentially, indicating that temporal context associations were present.

**Figure 1–Figure supplement 2.**
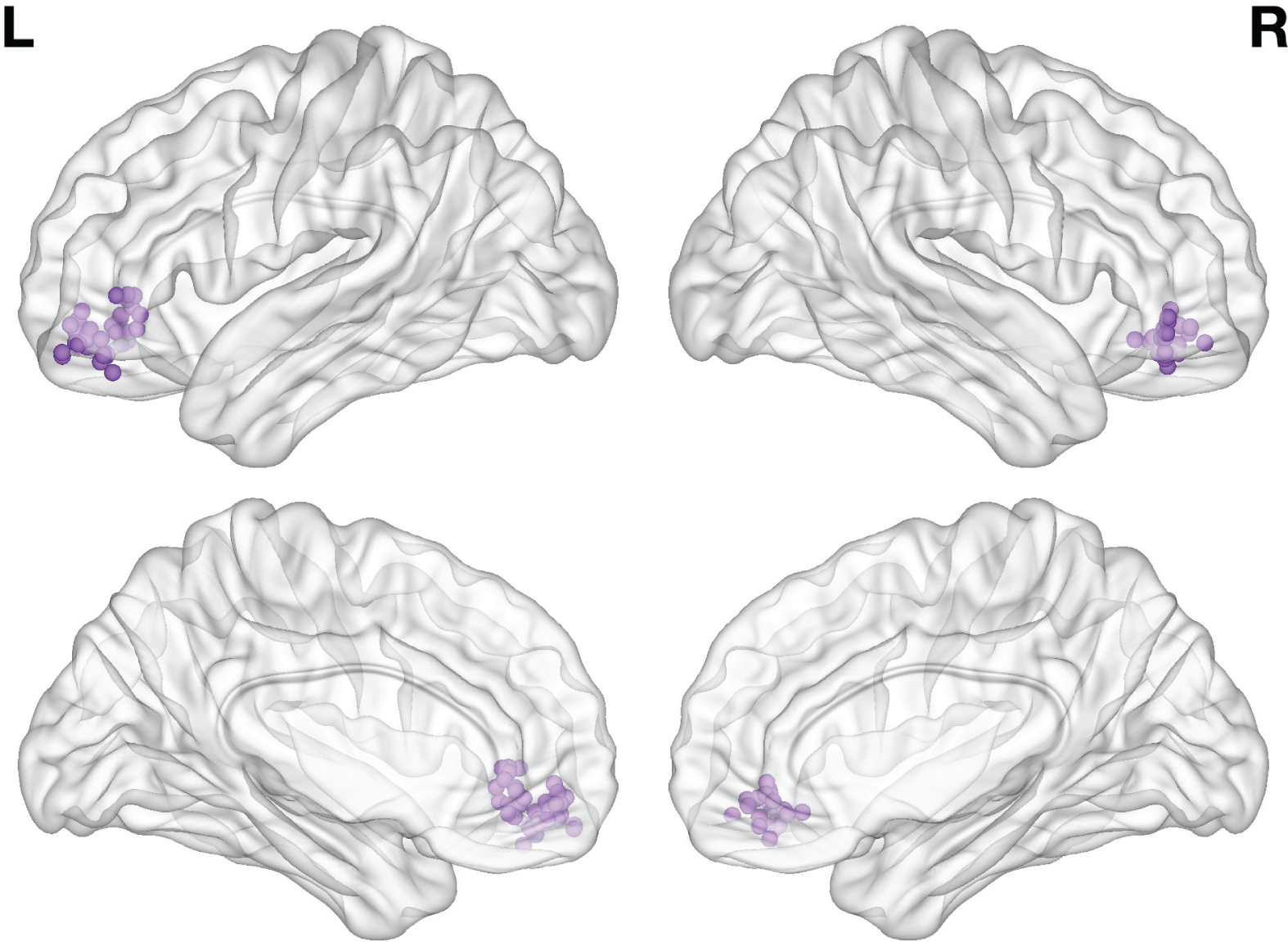
Electrode MNI locations for bilateral aVLPFC electrodes. Each sphere represents a single electrode.

**Figure 2–Figure supplement 1.**
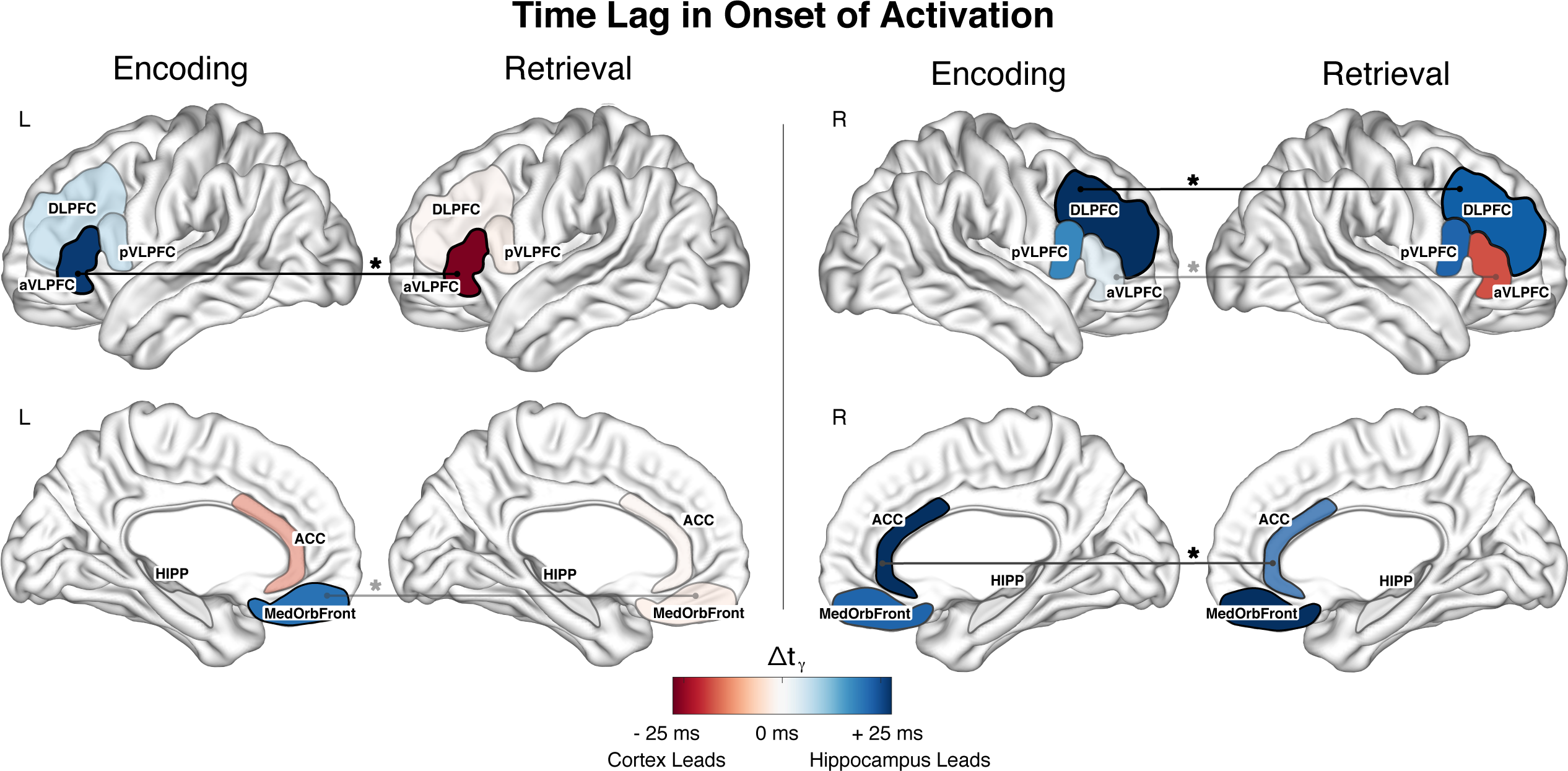
Mean Δ*t*_*γ*_ across electrodes for all PFC regions using a threshold of 2 times the calculated threshold. The left aVLPFC shows a Δ*t*_*γ*_ reversal between conditions that is consistent with the result using the original threshold. The magnitude of the Δ*t*_*γ*_ is larger for the higher threshold (mean encoding lag is +21.1 msec and mean retrieval lag is −18.9 msec). The left DLPFC also shows results that are consistent with the original threshold, with no Δ*t*_*γ*_ reversal between conditions and a significantly longer Δ*t*_*γ*_ during encoding versus retrieval.

**Figure 3–Figure supplement 1.**
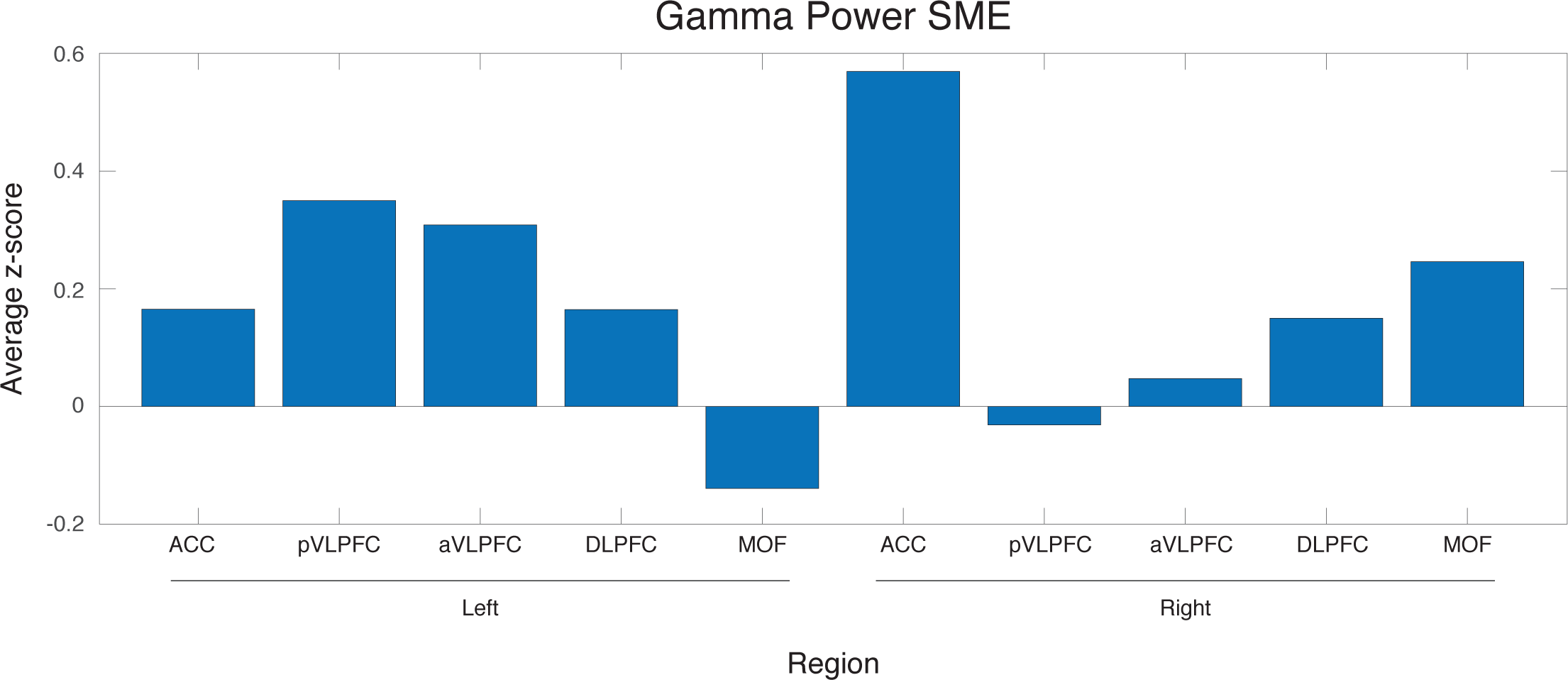
Subsequent memory effect in gamma band power for all regions. Z-scores were calculated for each region using a paired t-test between gamma power for recalled words and non-recalled words. A positive z-score indicates that the non-recalled gamma band power is greater than recalled gamma band power.

